# The proton-activated chloride channel inhibits SARS-CoV-2 spike protein-mediated viral entry through the endosomal pathway

**DOI:** 10.1101/2025.03.12.642872

**Authors:** Nicholas Koylass, Jaiprasath Sachithanandham, James Osei-Owusu, Kevin Hong Chen, Henry Yi Cheng, Andrew Pekosz, Zhaozhu Qiu

## Abstract

SARS-CoV-2 binds to its obligatory receptor, angiotensin-converting enzyme 2 (ACE2) and capitalizes on decreasing endosomal acidity and cathepsin-mediated spike protein cleavage to enter cells. Endosomal acidification is driven by V-ATPase which pumps protons (H^+^) into the lumen. The driving force for H^+^ is maintained by the import of chloride (Cl^-^) which is mediated by intracellular CLC transporters. We have recently identified the Proton-Activated Chloride (PAC) channel as a negative regulator of endosomal acidification. PAC responds to low pH and releases Cl^-^ from the lumen to prevent endosomal hyperacidification. However, its role in SARS-CoV-2 viral entry remains unexplored. Here, we show that overexpressing the PAC channel in ACE2 expressing HEK 293T cells markedly inhibited the SARS-CoV-2 spike-mediated viral entry. Several lines of evidence suggest that this effect was due to the suppression of the endosomal entry pathway. First, the abilities of PAC to regulate endosomal acidification and inhibit pseudoviral entry were both dependent on its endosomal localization and channel activity. Second, the inhibitory effect on viral entry was similar to the suppression mediated by E64-d, a cathepsin inhibitor, while no major additive effect for both treatments was observed. Third, this inhibition was also attenuated in cells expressing TMPRSS2, which provides the alternative entry pathway through cell surface. Importantly, PAC overexpression also inhibited the number and size of plaques formed by two live SARS-CoV-2 isolates (B.1 and Omicron XBB.1.16) in Vero E6 cells. Altogether, our data indicates that PAC plays a vital role in inhibiting SARS-CoV-2 viral entry and identifies this endosomal channel as a potential novel target against the infection of SARS-CoV-2 and other viruses, which rely on the endosomal pathway.

## 1 Introduction

Severe acute respiratory syndrome coronavirus 2 (SARS-CoV-2) is a positive sense RNA coronavirus responsible for the COVID-19 pandemic. mRNA vaccines and the induction of virus neutralizing antibody responses are the primary prophylactic treatment against severe COVID-19 ^1^. However, as the virus continues to mutate to evade preexisting population immunity, the efficacy of current vaccines wanes ^2^. Identifying novel treatments and prophylactics that target viral entry but are independent of the mutations that drive vaccine immune escape would be extremely useful in the evolutionary arms race against SARS-CoV-2.

SARS-CoV-2 binds to its major receptor, the angiotensin converting enzyme 2 (ACE2), via the receptor binding domain that is present in its spike protein. The spike protein is divided into two domains with the S1 domain mediating receptor binding and the S2 domain mediating cell membrane fusion ^3^. The S1/S2 boundary is cleaved by furin, a Ca^2+^-dependent serine endopeptidases, exposing the S2’ site ^4,5^. The virus enters cells through clathrin-mediated endocytosis ^6^ and the exposed S2’ site is cleaved by pH-dependent cysteine proteases such as Cathepsins B/L in the endolyosomal pathway ^7^. Previous work showed that disrupting the H^+^ gradient with lysomotropic agents such as bafilomycin and chloroquine ^8^ or inhibiting cathepsins, with E64-d reduces SARS-CoV-2 viral entry ^7,9^. Alternatively, S2’ cleavage can occur at or near the cell surface by another protease TMPRSS2 ^10,11^. The TMPRSS2 pathway is thought to be the primary mode of viral entry for early strains of SARS-CoV-2 such as B.1 which contains the D614G mutation, and the Delta variant ^12,13^. Conversely, Omicron variants prefer to use the endosomal path to enter cell ^14–16^.

During acidification, the V-ATPase pumps proton into the endosomal lumen ^14,15^. H^+^ accumulation increases the membrane potential which requires a counter-ion to maintain electroneutrality. Endosomal CLC exchangers (CLC3–5) fulfill this role by transporting one H^+^ out and two Cl^-^ into the lumen ^17,18^. This continual exchange maintains the driving force for H^+^. The accumulation of both H^+^ and Cl^-^ mediates endosomal acidification. Recently, we have identified a new player, the proton-activated Cl^-^ (PAC) channel that localizes to endosomes^19^. PAC is activated by decreasing endosomal pH and mediates the efflux of Cl^-^ out of endosomes ^19^. As such, it acts as a brake to antagonize endosomal acidification ^19^. Since SARS-CoV-2 requires acidic endosomal pH to enter cells, we hypothesized that overexpressing PAC would reduce SARS-CoV-2 viral entry by increasing endosomal pH.

Here, using a pseudoviral luciferase assay, we show that overexpressing PAC inhibits SARS-CoV-2 spike-mediated viral entry by preventing endosomal acidification. Using a plaque assay, we further find that PAC overexpression reduces viral infection of live SARS-CoV-2 strains B.1 (D614G) and XBB.1.16 (Omicron) in Vero E6 cells. Our study establishes this newly identified endosomal chloride channel as a potential novel target against SARS-CoV-2 infection.

## 2 Materials & Methods

### 2.1 Cell Maintenance

Tetracycline inducible HEK 293T cells (T-REx) were maintained in Dulbecco’s modified Eagle’s medium (DMEM) supplemented with 10% tetracycline-free fetal bovine serum (FBS) and 1% penicillin/streptomycin (P/S) and 1% Glutamax™ Supplement at 37°C in humidified 95% CO_2_ incubator. HEK 293T cells and Vero E6 cells were grown under the same conditions but with FBS that contained trace amounts of tetracyline.

### 2.2 Cloning and mutagenesis

FLAG-tagged human PAC mutants (Y10A/L13A and L309C) were generated from the previously designed human PAC (pcDNA5/FRT/TO) plasmid ^19^. The QuikChange II XL site-directed mutagenesis kit (Agilent Technologies) was used to generate Y10A/L13A and L309C constructs. Each construct was co-transfected with FLP recombinase (pOG44, Life technologies) into Flp-In HEK 293T cells. Cells were selected with 200 ug/ml hygromycin and single cell clones were identified.

To generate the PAC-mCherry fusion plasmid (PAC-mCherry), we used restriction enzyme cloning to generate the PAC-mCherry fusion plasmid by inserting a Gly-Ser-Gly-Ser linker between PAC and mCherry cDNA sequences

### 2.3 Production of lentiviral particles

HEK 293T cells were seeded at 70% confluency into 10 cm dishes 24 hours prior to packaging lentiviral particles expressing ACE2, TMPRSS2 or hPAC-mCherry. 6 μg of ACE2 plasmid (pLENTI_ACE2_PURO, Addgene 155295), TMPRSS2 plasmid (pLEX307-TMPRSS2-blast, Addgene158458) or hPAC-mCherry were co-transfected with 2 μg of packaging plasmids [pVSV-G (pMD2.G, Addgene 12259), pMDL (pMDLg/pRRE, Addgene 12251) and pRSV (pRSV-REV, Addgene 12253)].

To generate VSV-G pseudoviral particles, we transfected a 15 cm dish of HEK 293T cells with 3.5 μg of VSV-G along with 3.5 μg of packaging plasmids pMDL and pRSV, and 16 μg of luciferase construct (pHIV-Luc-ZsGreen, Addgene 39196).

To generate SARS-CoV and SARS-CoV-2 pseudoviral particles, we transfected a 15 cm dish of HEK 293T cells with 5.4 μg of SARS-CoV or SARS-CoV-2 plasmid (see the SARS-CoV and list of SARS-CoV-2 pseudoviruses in Supplementary Table 1) along with 3.5 μg of packaging plasmids pMDL and pRSV, and 16 μg of the luciferase construct.

All transfected cells were incubated at 37 °C with 5% CO_2_ for 18 hours before the culture media was switched with fresh culture media containing 1% BSA. Viral supernatants were collected 48 hours post-transfection, centrifuged, filtered through a 0.45 μm PES filter (GE Healthcare, cat. 67802504), aliquoted, and stored at -80°C.

### 2.4 Generation of stably expressing cells

To generate ACE2-expressing cells, HEK 293T T-REx cells were infected with ACE2 lentiviruses in a 6-well plate along with 8 μg/ml polybrene for 48 hours at 37°C to assist with the transduction and then drug selected with 2 μg/ml puromycin for 48 hours. TMPRSS2-expressing cells were generated in a similar manner but without drug selection.

To generate PAC-overexpressing cells, Vero E6 cells were infected with PAC-mCherry lentiviruses in a 6-well plate along with 8 μg/ml polybrene. Lentiviruses were spinfected onto Vero E6 cells for 30 minutes at 3000 g at 4°C. Cells were then incubated for 48 hours at 37°C to ensure optimal transduction.

### 2.5 Pseudoviral Luciferase Assay

HEK 293T T-REx ACE2-expressing cells were seeded onto 100 μM poly-L-lysine coated 96 well plates and treated with 10 ng/ml tetracycline to induce PAC expression for 24 hours before infection. Cells were then treated with fresh media with pseudoviral particles and tetracycline for 48 hours before lysed with Bright Glo (Promega) luciferase reagent. Lysates were transferred to 96 well opaque white plates (Greiner 655083) and the luciferase signal was acquired using the Infinite M plex Tecan plate reader over a 1-minute integration time.

When assessing the role of cathepsins in viral entry, cells were pretreated for two hours with 10 μM E64-d (Alostatin) before adding pseudoviruses and tetracycline. E64-d was maintained for the duration of the pseudoviral infection.

### 2.6 Western blot

Cells were lysed with RIPA buffer (Sigma) containing a cOmplete™, Mini, EDTA-free Protease Inhibitor Cocktail (Roche) and centrifuged at 20,000 g for 10 min in 4°C. Proteins were separated using 4–20% Novex™ Tris-Glycine Mini Protein Gels (Thermo Fischer Scientific) and NuPAGE MES SDS Running Buffer and transferred onto a nitrocellulose membrane. The membrane was then blocked with 5% Blotting Grade Blocker Non-Fat Dry Milk (Bio-Rad) in tris-buffered saline, with tween-20 (TBST) for 30 min at room temperature, washed three times and incubated with primary antibodies: mouse anti-PAC (1:200; in house), rat anti-ACE2 (1:5000; Biolegend 375801; CloneA200691), mouse anti-GAPDH (1:1000; CST 97166) or rabbit anti-TMPRSS2 (1:5000; abcam ab109131; Clone EPR3862) at 4°C for overnight. Primary antibodies were diluted in 5% BSA and 0.02% sodium azide and secondary antibodies were diluted in 5% Blotting Grade Blocker Non-Fat Dry Milk. Membranes were washed three times with TBST and then incubated at room temperature with the corresponding Amersham ECL horseradish peroxidase (HRP)-conjugated secondary antibody (1:5000; Cytiva) for 1 hour at room temperature. Proteins were then detected using Pierce ECL Plus Western Blotting Substrate (Thermo Fisher Scientific) and images were prepared using ImageJ.

### 2.7 Immunofluorescence

Cells were fixed with 4% paraformaldehyde (PFA) followed by three phosphate-buffered saline (PBS) washes. Cells were then permeabilized with 0.1% Triton X-100 and incubated with primary antibodies (mouse anti-FLAG (1:1000; Origene TA50011) and rabbit anti-EEA1 (1:1000; CST C45B10) overnight at 4°C in PBS containing 3% BSA. Cells were then stained with Alexa Fluor-conjugated secondary antibodies (1:1000; Thermo Fisher Scientific) incubation for 1 hour at room temperature. Coverslips were then mounted on microscope slides and images were taken using Zeiss LSM900 confocal microscope.

### 2.8 Electrophysiology

Whole-cell patch clamp recordings were performed as described previously ^20^. Cells were plated on coverslips and PAC expression was induced with 10 ng/ml tetracycline 24 hours before recordings. Cells were recorded in an extracellular solution (ECS) containing (in mM): 145 NaCl, 1.5 CaCl_2_, 2 MgCl_2_, 2 KCl, 10 HEPES, and 10 glucose. pH was adjusted to pH 7.3 with NaOH and osmolality was 297–310 mOsm/kg. Acidic ECS of pH 4.6 or pH 5.0 was made with the same ionic composition but with 5 mM Na_3_-citrate buffer. ECS was applied using a local gravity perfusion system. Recording pipettes (2–7MΩ) were filled with internal solution containing (in mM): 135 CsCl, 2 CaCl_2_, 1 MgCl_2_, 4 MgATP, 0.5 Na_3_ -GTP, and 5 EGTA (pH adjusted to 7.2 with CsOH and osmolality was 280–290 mOsm/kg). Recordings were done at room temperature with MultiClamp 700B amplifier and 1550B digitizer (Molecular Devices). Data acquisition was performed with pClamp 10.7 software (Molecular Device), filtered at 2 kHz and digitized at 10 kHz. Voltage ramp pulses were applied every 5 s from −100 to +100 mV at a holding potential of 0 mV.

### 2.9 Endosomal pH measurement

Endosomal pH was measured by a ratiometric method as described previously ^21^. Cells were plated onto poly-L-lysine coated plates and PAC expression was induced 24 hours prior to the experiments. Cells were starved in serum-free media for 30 min to remove residual transferrin and incubated with 75 μg/mL pH-sensitive FITC-transferrin and 25 μg/mL pH insensitive Alexa Fluor 633-transferrin (both from Thermo Fisher Scientific) at 37°C for 1 hour. The transferrin uptake was quenched by placing cells on ice. Cells were then washed once with ice-cold neutral PBS to remove excess transferrin, followed by a PBS wash at pH 5.0 to remove surface bound transferrin and a final wash in neutral PBS. Cells were trypsinized and analyzed by flow cytometry. FITC-transferrin and Alexa Fluor 633-transferrin double positive cells were gated and a ratiometric analysis of ∼30,000 cells for each biological replicate was used to determine the endosomal pH. A standard curve was constructed using the Intracellular pH Calibration Buffer kit (Thermo Fisher Scientific). The potassium ionophore valinomycin and the potassium/hydrogen ionophore nigericin were used to equilibrate the endosomal pH with the calibration buffers at a concentration of 10 μM.

### 2.10 Plaque Assay

SARS-CoV-2 isolates representative of an early, spike D614G lineage (SARS-CoV-2/USA/DC-HP00007/2020, GISAID sequence EPI_ISL_434688; Pango lineage B.1) and Omicron variant XBB1.16 lineage (hCoV-19/USA/MD-HP46342-PIDFPPISFA/2023, GISAID sequence EPI_ISL_ 17394984) were serially diluted and added to wells containing either Vero E6 or Vero E6 PAC expressing cells. Wells were incubated for 1 hour at 37°C before an agar/media mixture was carefully added to the wells. The virus was incubated for 60 hours. Wells were fixed with 4% formaldehyde overnight at room temperature and stained with naphthol blue black for another 24 hours before counting plaques ^22^. Plaque areas were determined by imaging wells and a standard ruler using a Nikon Fluorescence Dissection Microscope with an Olympus DP-70 camera. Images of wells were transferred to ImageJ software and a pixel-to-mm scale was established using the ruler. Plaques were traced and area (in mm^2^) was determined based on selected pixels ^23^.

### 2.11 Quantification, statistical analysis and figure preparation

Data and statistical analyses were performed using Excel, Clampfit 10.7, ImageJ and GraphPad Prism 10. Statistical analysis between two groups were performed using a two-tailed Student’s *t*-test with Welch’s correction. Multiple group comparisons were performed using one-way analysis of variance (ANOVA) with a Bonferroni post hoc test or two-way ANOVA using the Sidak test. Non-significant ‘ns’ *p* values were not reported. All numeric data are shown as mean ± SEM and the significance level was set at *p < 0.05*. Figures were prepared in Inkscape, and models were prepared in Biorender.

## 3 Results & Discussion

To investigate the role of PAC in SARS-CoV-2 spike protein-mediated entry, we stably expressed ACE2 in HEK 293T cells without (control) or with tetracycline-induced PAC expression. ACE2 expression and PAC induction were validated via western blot (Supplemental Figure 1A). The increased PAC channel activity in tetracycline-induced cells was further confirmed using whole cell patch clamp electrophysiology (Supplementary Figure 1B and 1C). Cells were infected with pseudoviral particles that co-expressed various envelope proteins along with the firefly luciferase reporter gene for quantifying viral entry. While tetracycline treatment had no effect on SARS-CoV-2 spike-mediated pseudoviral entry in control cells, PAC overexpression robustly decreased the viral entry mediated by the spike protein of SARS CoV-2, but not the negative control vesicular stomatitis virus G glycoprotein (VSV-G) (Figure 1A). SARS-CoV also binds ACE2 and enters cells via the endosomal pathway. Similar to SARS CoV-2, PAC overexpression also inhibited SARS-CoV spike-mediated pseudoviral entry. These data suggest that the PAC channel may play a broad role in cell entry of viruses that depend on the endosomal pathway. We expanded the study and tested the spike proteins of other major SARS-CoV-2 variants. Indeed, we found that overexpression of PAC reduced all of their cell entries (Figure 1B).

**Figure 1.**
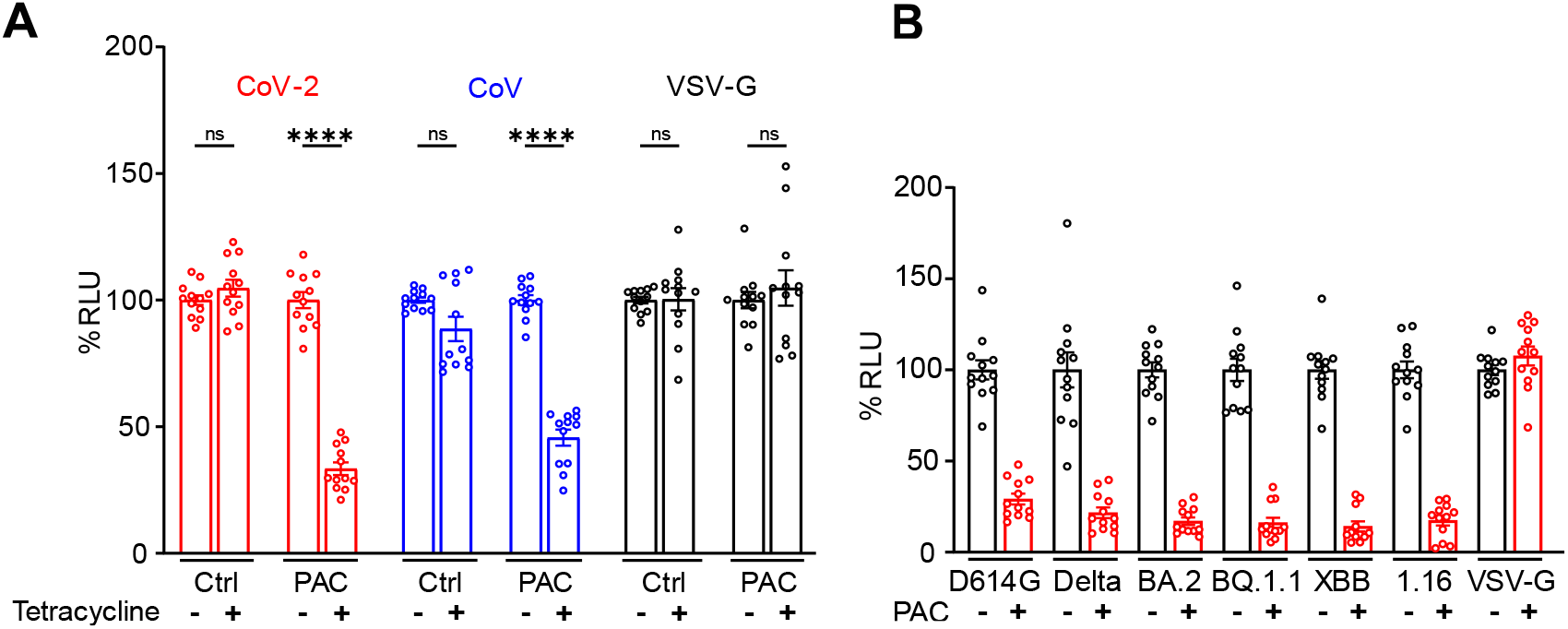
PAC overexpression inhibits SARS-CoV1/2 spike-mediated pseudoviral entry. A) HEK 293T ACE2 cells (CTRL) or PAC inducible cells (PAC) treated with SARS-CoV-2, SARS CoV or VSV-G pseudovirus. % Relative Luciferase Units (RLU) were normalized to the mean of the no-tetracycline controls. n = 12 wells from 4 experiments, two-way ANOVA with multiple comparisons. B) PAC inducible HEK 293T ACE2 cells were infected with pseudoviruses carrying different SARS-CoV-2 spike proteins. SARS-CoV-2 XBB.1.16 is labelled 1.16 in figure. n = 12 wells from 4 experiments, unpaired two-tailed student’s *t-*test with Welch’s correction. The % RLU between control and tetracycline induced conditions were statistically significant with *p <0.0001* for all groups except VSV-G pseudoviruses. Bars represent mean ± SEM; ns = not significant, *\*\*\*\* p < 0.0001*.

We next probed the mechanism whereby PAC impedes SARS-CoV-2 spike-mediated viral entry. We first measured endosomal pH in HEK 293T ACE2 cells using a ratiometric transferrin assay ^19^. Consistent with our previous findings ^19^, PAC induction resulted in a hypo-acidified endosomal pH, by ∼0.4 pH units, compared to that of control cells (Figure 2A). Since SARS-CoV-2 and SARS-CoV require proper endosomal acidification, the increased endosomal pH in PAC-expressing cells likely explain the decreased viral entry. Human PAC traffics to endosomes via the classic YxxL motif where Y is tyrosine, x is any amino acid and L is leucine ^19^.

**Figure 2.**
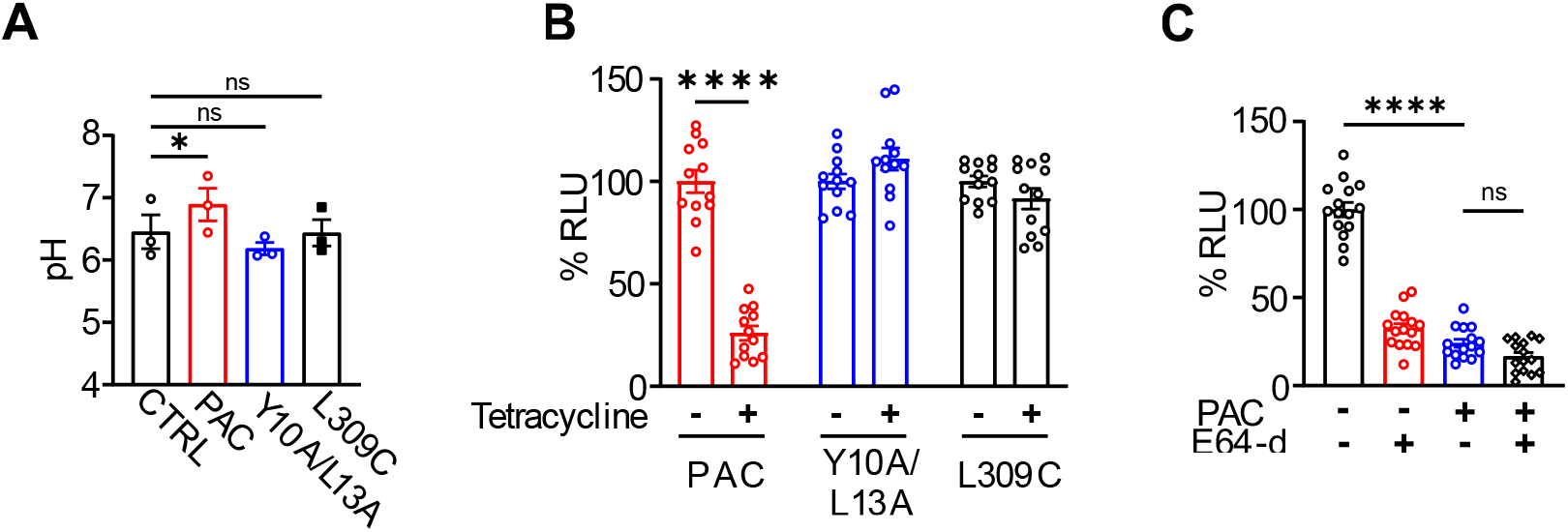
The inhibitory effect on SARS-CoV-2 spike-mediated pseudoviral entry by PAC depends on its endosomal localization and channel activity. A) Endosomal pH measurement using ratiometric transferrin assay for control HEK 293T ACE2 cells and the inducible cells expressing PAC, Y10A/L13A, and L309C mutants. n = 3, one-Way ANOVA with multiple comparisons. B) SARS-CoV-2 spike-mediated pseudoviral entry for cells with or without expression of WT PAC, Y10A/L13A, and L309C mutants. % Relative Luciferase Units (RLU) were normalized to the mean of the no-tetracycline controls. n = 12 wells from 4 experiments, two-way ANOVA with multiple comparisons. C) Cathepsin inhibitor E64-d had no major additive effect on SARS-CoV-2 spike-mediated pseudoviral entry with PAC overexpression. % RLUs were normalized to the mean of the non-treated control. n = 15 wells from 5 experiments, one Way ANOVA with multiple comparisons. Bars represent mean ± SEM; ns = not significant, *\* p<0.05, **** p < 0.0001*.

We previously generated single trafficking mutants (Y10A and L13A) and showed that the ability of PAC to regulate endosomal pH is dependent on its localization to endosomes ^19^. Here, we generated a double mutant (Y10A/L13A) and validated its protein expression and predominant cell surface localization by western blot and immunostaining, respectively (Supplementary Figure 2A and 2B). In contrast with WT PAC, cells expressing the Y10A/L13A trafficking mutant did not exhibit endosomal hypo-acidification, indicating that the endosomal localization is required for the regulation of endosomal pH by PAC (Figure 2A). Interestingly, these cells also failed to inhibit SARS-CoV-2 spike-mediated pseudoviral entry (Figure 2B). To further test the involvement of PAC channel activity, we generated HEK 293T cells with inducible expression of L309, a channel “dead” mutant, whose expression was validated by western blot (Supplemental Figure 2A) and by whole cell patch clamp electrophysiology ^20^ (Supplementary Figure 2C and 2D). Unlike WT PAC-expressing cells, L309C-expressing cells had normal endosomal pH and did not suppress SARS-CoV-2 spike-mediated pseudoviral entry (Figure 2 A and 2B). Together, these data suggest that endosomal localization and channel activity of PAC are both required for its regulation of endosomal pH, which in turn is critical for SARS-CoV-2 spike-mediated viral entry.

When SARS-CoV-2 binds to ACE2, the ACE2/virus particle complex enters the cell via the endocytic pathway. The S2′ site on the spike is then cleaved by cathepsins B/L ^7,24^, which are pH-dependent cysteine proteases in endosomes ^7^. This key event releases the fusion peptide and initiates fusion pore formation ^11^. The cathepsin inhibitors, such as E64-d (Aloxistatin), are known to impair SARS-CoV-2 viral entry ^7,9^. Indeed, we found that E64-d treatment inhibited SARS-CoV-2 spike-mediated pseudoviral entry (Figure 2C). Importantly, treating PAC-expressing cells with E64-d did not further generate any compounding effect on SARS-CoV-2 spike-mediated pseudoviral entry. This result suggests that PAC and cathepsins may regulate viral entry through affecting the same endosomal pathway. It is therefore consistent with our hypothesis that PAC overexpression blocks SARS-CoV-2 viral entry by hindering endosomal acidification, which is required for pH-dependent cathepsin activities.

SARS-CoV-2 can also utilize trypsin-like proteases, such as TMPRSS2, on the cell surface to directly enter cells without endocytosis ^12^. Endosomal alkalizers such as chloroquine and hydroxychloroquine fail to inhibit SARS-CoV-2 viral entry in cells with high TMPRSS2 expression ^25^, suggesting that TMPRSS2 may dictate the route how SARS-CoV-2 infect cells ^12^. HEK 293T ACE2 cells used in the above pseudoviral entry assay do not express endogenous TMPRSS2. To test if PAC regulates viral entry through the endolyosomal pathway, we stably expressed TMPRSS2 in the PAC-inducible HEK 293T cells (Figure 3A) and performed the SARS-CoV-2 pseudovirus assay. Although PAC overexpression still reduced SARS-CoV-2 pseudoviral entry in the presence of TMPRSS2, the inhibitory effect was attenuated compared to the cells without TMPRSS2 expression (Figure 3B). Together, these data further suggest that PAC suppresses SARS-CoV-2 spike-mediated pseudoviral entry through interfering with endosomal acidification and blocking the endosomal viral entry pathway.

**Figure 3.**
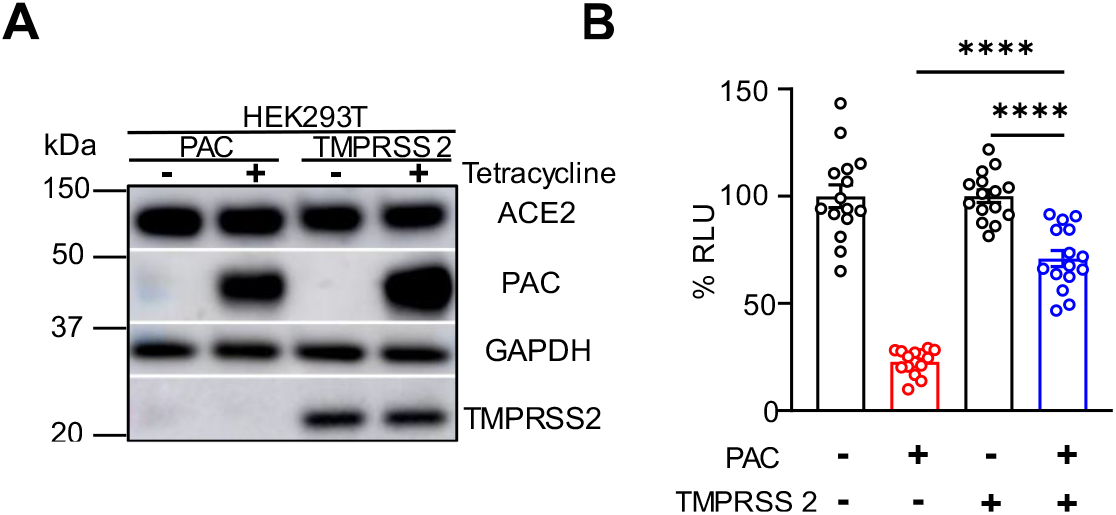
TMPRSS2 expression attenuates the inhibitory effect of PAC on SARS-CoV2 spike-mediated pseudoviral entry. A) Western blot validating the inducible expression of PAC and stable expression of ACE2 and TMPRSS2. Representative image of three experiments. B) SARS-CoV-2 spike-mediated pseudoviral entry for HEK 292T ACE2 cells with or without PAC and TMPRSS2 expression. % Relative Luciferase Units (RLU) were normalized to the mean of the controls that did not express PAC or TMPRSS2. n = 15 wells from 5 experiments, unpaired two-tailed student’s *t-*test with Welch’s correction. Bars represent mean ± SEM; **** *p<0.0001*.

Pseudoviruses are replication incompetent viral particles that contain backbone and surface proteins from different viruses ^26^. These viruses were widely used during the COVID-19 pandemic, to investigate how SARS-CoV-2 recognizes host cell receptors, its mechanism of entry and aided in the rapid development of testing vaccines ^26^. However, because pseudoviruses are devoid of pathogenic genes ^36^, they are unable to provide insight into viral replication, release or transmission ^27^. Therefore, we sought to further test the effect of PAC overexpression on live SARS-CoV-2 in Vero E6 cells ^28,29^. Vero E6 cells were derived from the kidney of an African green monkey which express endogenous ACE2 receptor ^30^. They are suitable for studying viral entry via the endosomal pathway since they do not express TMPRSS2 ^31,32^.

We first stably expressed PAC in Vero E6 cells and assessed its expression by Western blot (Supplemental Figure 3A). Next, we performed whole cell patch clamp electrophysiology and verified the increased PAC channel activity (Supplementary Figure 3B and 3C). The plaque assay is the gold standard assay, used to count discrete infectious centers ^28,29^. The infectious centers are used to assess viral replication and infectivity as plaque forming units (PFU/ml). We performed the plaque assay using two distinct strains of SARS-CoV-2: B.1 (D614G) and XBB.1.16 (Omicron). The B.1 strain is one of the earliest versions of SARS-CoV-2 that contains the D614G mutation. We used a SARS-CoV-2 spike protein that only contained the D614G mutation for all experiments (except for Figure 1C) of our pseudoviral experiments ^33^. Interestingly, we observed a 5.4-fold decrease in plaque forming units in PAC-expressing Vero E6 cells compared to control cells (Figure 4A and 4B). To test the effect of PAC overexpression on viral infection of other SARS-CoV-2 strains. We chose XBB.1.16, a sub-variant descended from Omicron BA.2. Again, we observed a similar albeit more modest inhibition (2.5-fold decrease in plaque forming units) in PAC-expressing cells (Figure 4D and 4E). Compared to the B.1 strain, novel mutations in the Omicron strain that increase replication ^34^ and ACE2 binding ^35^ may be responsible for the reduced inhibitory effect. In addition to viral titer, we also analyzed the diameter of the plaques, which is a potential readout for the viral lytic capability, cell-to-cell spreading and viral release ^33,36^. Interestingly, PAC overexpression also reduced plaque size of both B.1 and XBB.1.16 viral strains (Figure 4C and 4F). Together, these results are consistent with the pseudoviral entry data, suggesting that PAC inhibits SARS-CoV-2 viral entry, and potentially viral replication as well.

**Figure 4.**
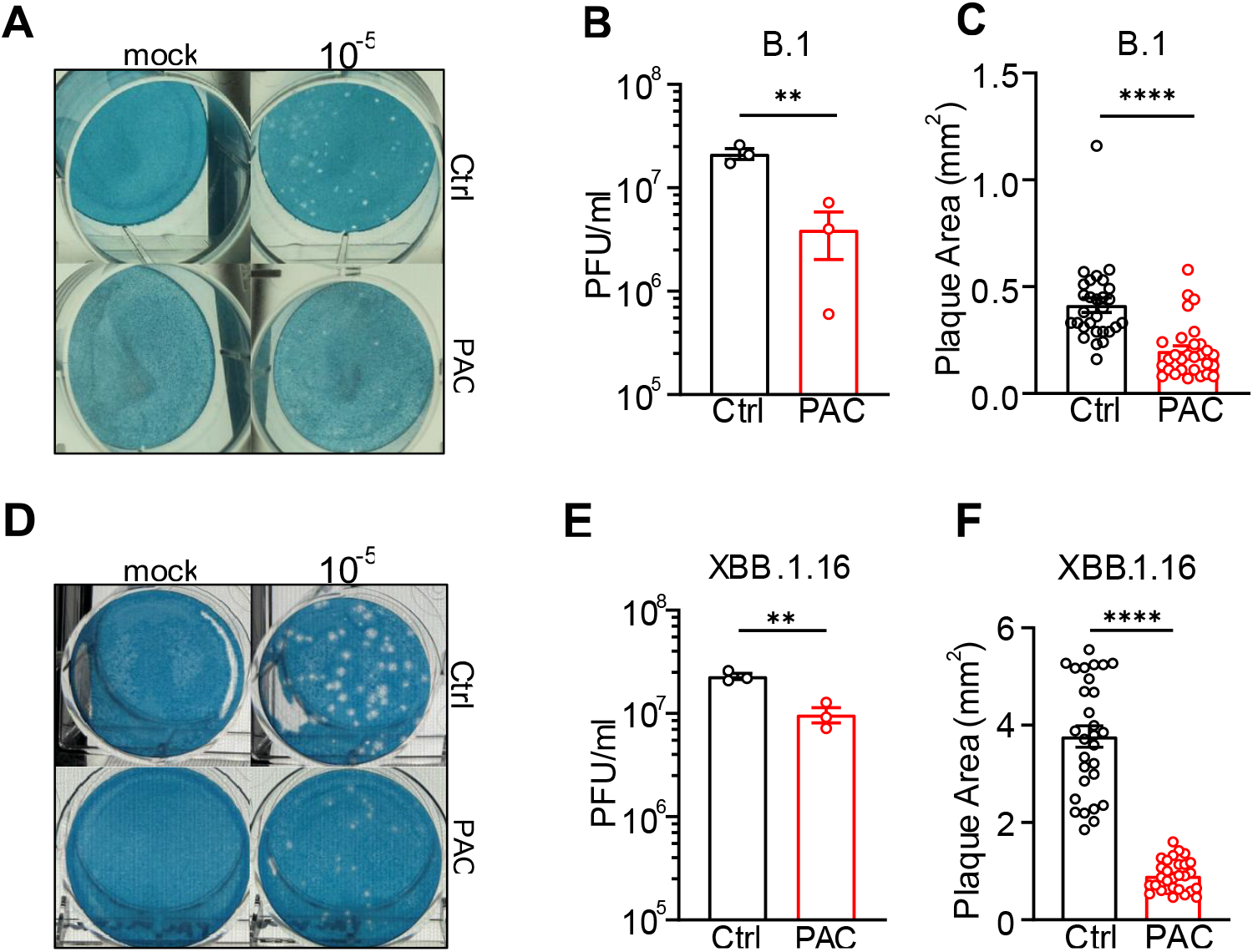
PAC overexpression inhibits viral infection of live SARS-CoV-2 strains in Vero E6 cells. A) Representative plaque assay, B) viral titer, and C) plaque size of SARS-CoV-2 B.1 virus. D) Representative plaque assay, E) viral titer, and F) plaque size of SARS-CoV-2 XBB.1.16 virus. n = 3 experiments for viral titer and n = 29 plaques for the size measurement, unpaired two-tailed student’s *t-* test with Welch’s correction. Bars represent mean ± SEM, *\*\* p<0.01, **** p<0.0001*.

In this study, we show that PAC overexpression prevented endosomal acidification, thereby suppressing spike protein-mediated viral entry of SARS-CoV and SARS-CoV-2. We further find that this inhibitory effect was diminished in TMPRSS2-expressing cells and did not produce additive effects together with the treatment of endosomal cathepsin inhibitor. Finally, PAC overexpression inhibited the replication and cytopathic effects of B.1 (D614G) and XBB.1.16 (Omicron) SARS-CoV-2 strains in Vero E6 cells. Based on these results, we present a model whereby overexpression of the PAC channel disrupts endosomal acidification, which then inhibits viral entry through the endosomal pathway (**Figure 5**).

**Figure 5.**
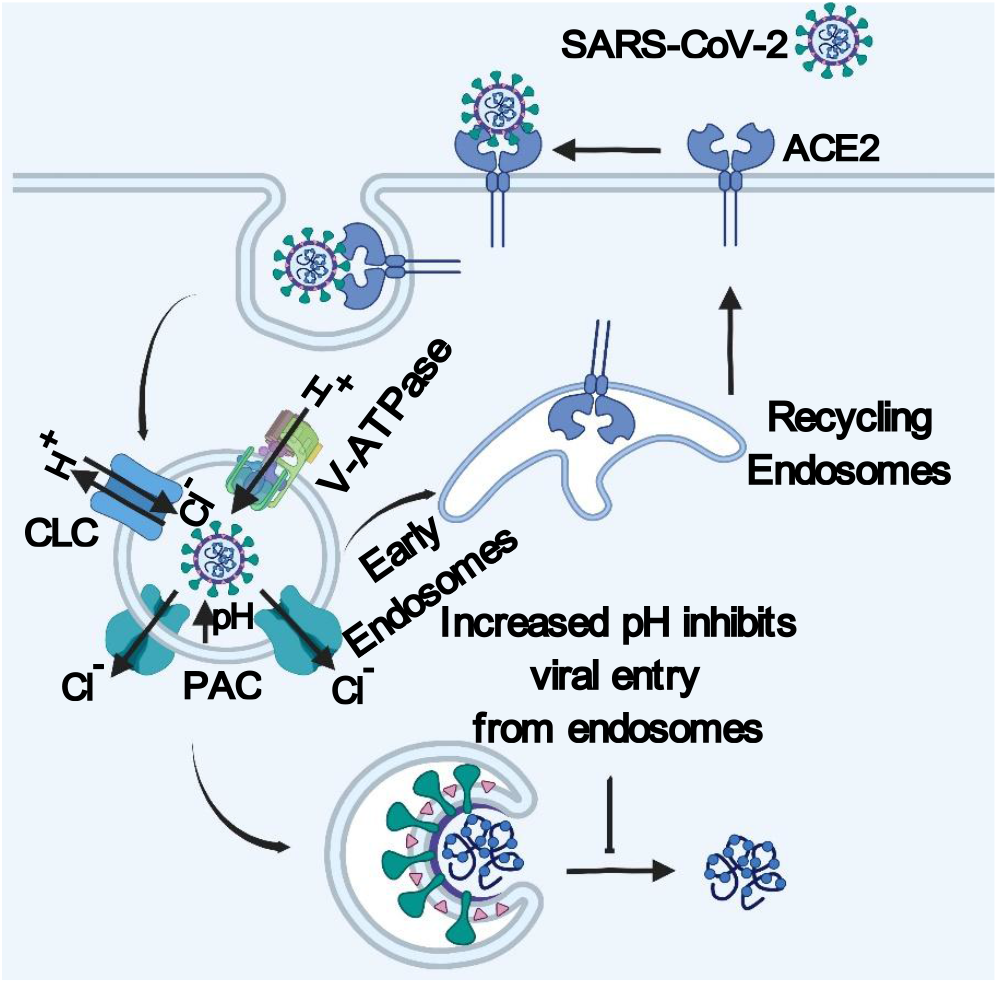
Model describing how PAC inhibits SARS-COV-2 viral entry. PAC overexpression increases endosomal pH thereby inhibiting SARS-CoV-2 viral entry.

Interestingly, a recent dCas9-based CRISPR activation screen in human lung epithelial Calu-3 cells has identified *TMEM206/PACC1*, the gene encoding the PAC channel, as one of the top hits with unknown mechanisms. Upregulation of PAC reduced the viral titer of SARS-CoV-2 by 2-3 logs ^39^. Our study on viral entry provides a potential molecular mechanism underlying this remarkable inhibitory effect. The relatively weaker effect on viral replication we observed here may be attributed to the differences in specific SARS-CoV-2 viral strain and cell types (Vero E6 vs. Calu-3) that were used in the studies. For example, the CRISPR screen study used the ancestral SARS-CoV-2 USA/WA-1/2020 ^39^. This strain does not contain mutations that give it a selective replicative advantage compared to the B.1 or Omicron strain that were used in our study.

Importantly, our work adds to the growing body of evidence that organelle ion channels are critical for SARS-CoV-2 viral entry and replication by regulating endosomal acidification ^40,41^. Recently, a study has reported that inhibiting or knocking down TRPM7 increases endosomal pH and negatively affects entry of enveloped viruses, such as SARS-CoV, SARS-CoV-2, and Ebola ^41^. TRPM7 is a nonselective for cation channel that localizes to endosomes and conducts an outward countercurrent of cations ^41^. This countercurrent relieves the build-up of positive charge within the endosomal lumen and favors the sustained H^+^ pumping activity of the V-ATPase ^42^. This ensures persistent endosomal acidification which is critical for viral entry ^41^. Similar to TRPM7, the PAC channel likely regulates viral entry of many other enveloped viruses that are dependent on efficient endosomal acidification.

Drugs that raise endosomal pH, such as chloroquine and hydroxychloroquine impair viral entry and viral replication ^8^. However, these drugs are unsuitable for clinical use primarily because of their unwanted side effects. Our findings suggest that the endosomal PAC channel could be a potential therapeutic target for COVID19. Future study is needed to identify specific PAC agonists to activate the channel and prevent endosomal acidification thereby potentially inhibiting viral entry and replication of SARS-CoV-2 viruses.

## Author contributions

N.K. conducted most experiments, designed the study, and analyzed the data. J.S. and A.P. performed the plaque assay and analyzed the data. J.O.O. generated the inducible cell lines. K.H.C. performed immunofluorescence experiments. H.Y.C. and N.K. performed pH measurements. Z.Q. supervised the project. N.K. and Z.Q. wrote the manuscript with input from all authors.

## Acknowledgements

We thank Junhua Yang and Minghua Fan for assistance with electrophysiology. This work was supported by grants from the National Institute of Health (R01NS118014, R35GM124824, and RF1NS134549) to Z.Q. Z.Q. was also supported by the American Heart Association (AHA) Established Investigator Award, McKnight Scholar Award, Klingenstein-Simon Scholar Award, Sloan Research Fellowship in Neuroscience, and Randall Reed Scholar Award.

## Data Availability Statement

The data that supports the findings of this study can be requested from the corresponding author.

