## Supplementary Files for "The proton-activated chloride channel inhibits SARS-CoV-2 spike protein-mediated viral entry through the endosomal pathway"

**Supplementary Figures**

**Supplementary Figure 1. Validation of inducible PAC expression and activity.** A) Western blot of tetracycline-induced PAC expression in HEK 293T ACE2 cells. Representative image of three experiments. B) Representative I/V curve and C) current densities of acid-induced PAC currents in the inducible HEK 293T ACE2 cells. n = 10 cells from Ctrl group. n = 8 cells from PAC overexpression group. Bars represents mean ± SEM.

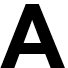

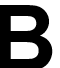

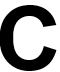

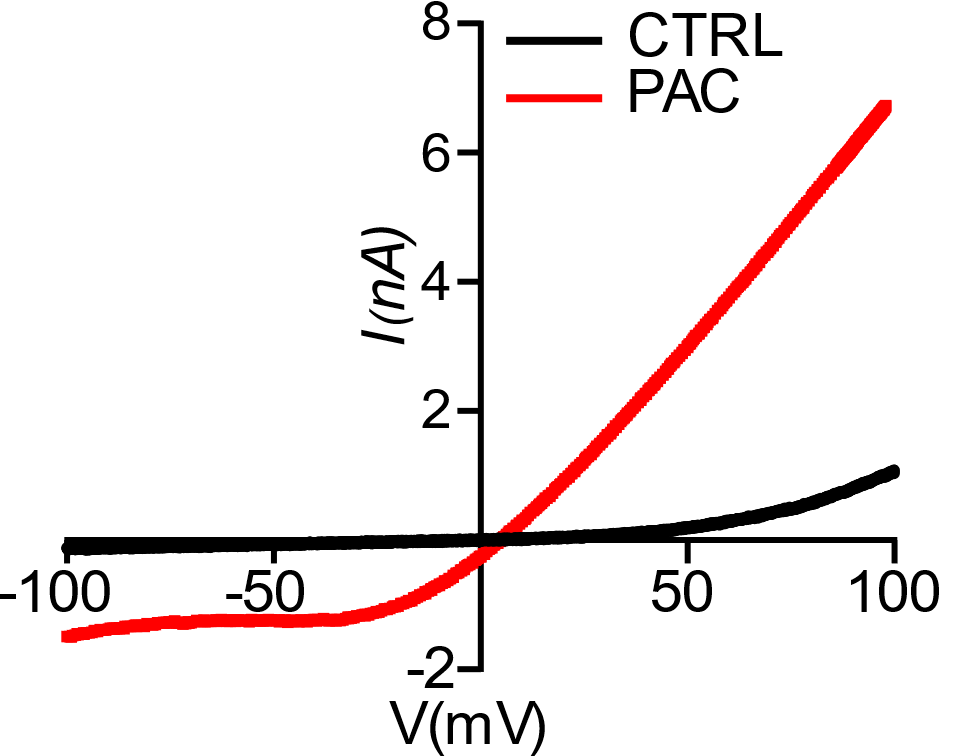

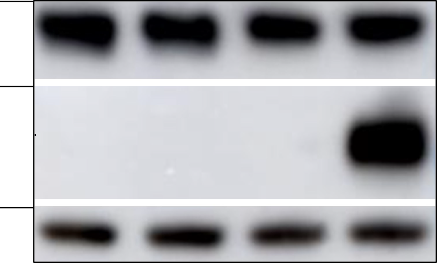

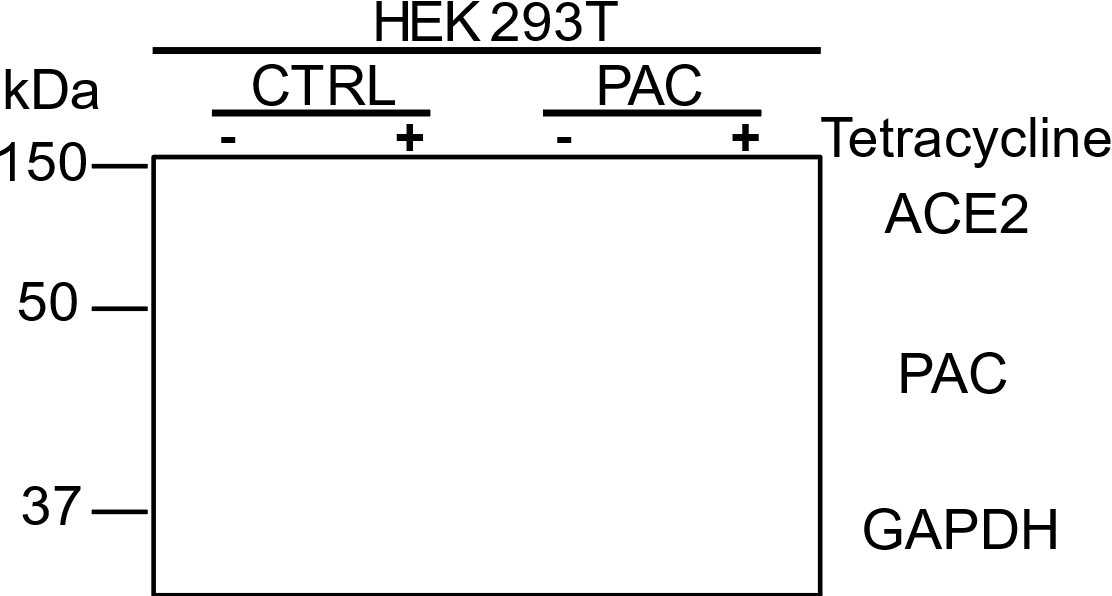

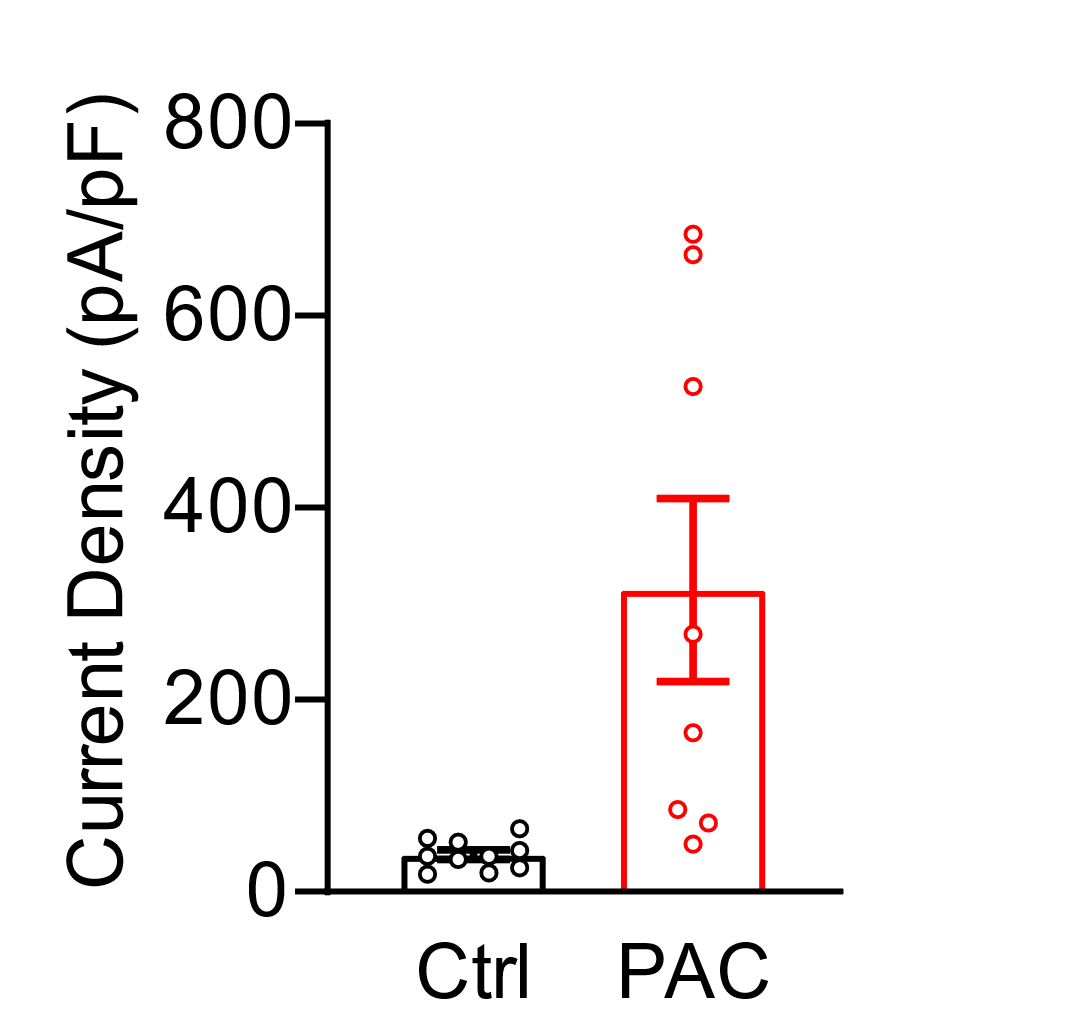

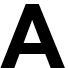

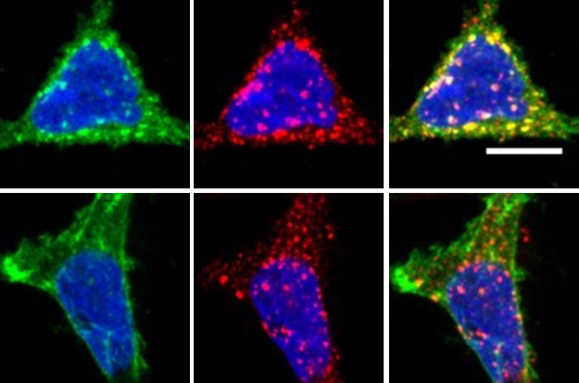

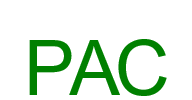

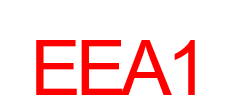

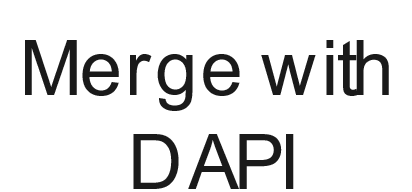

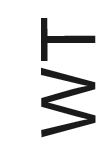

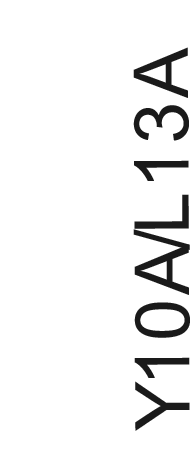

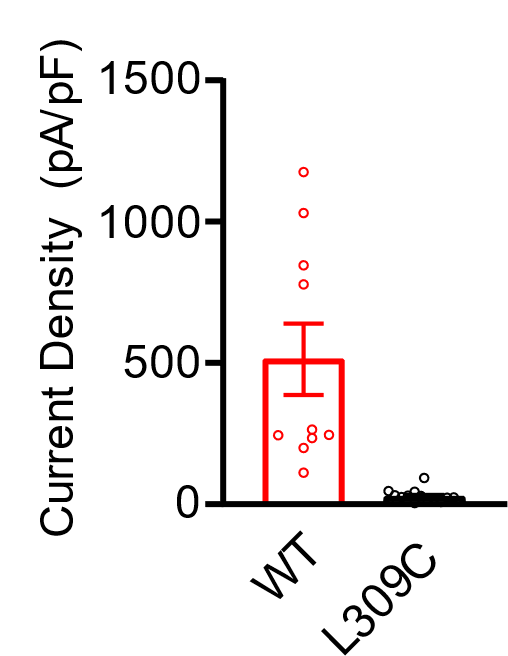

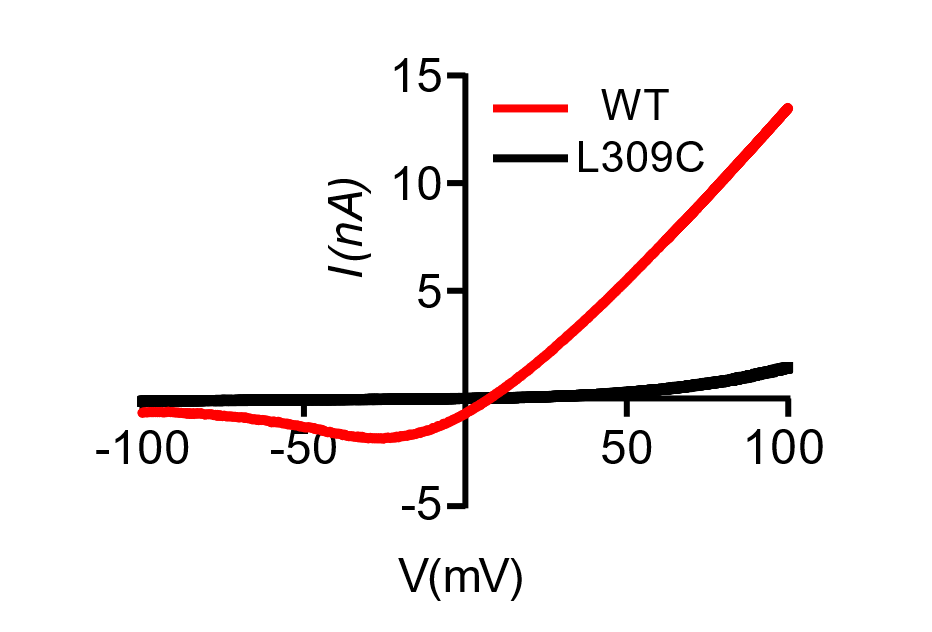

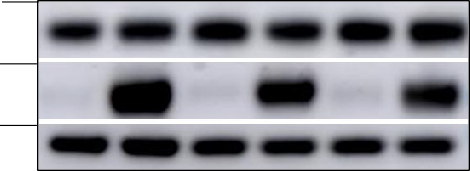

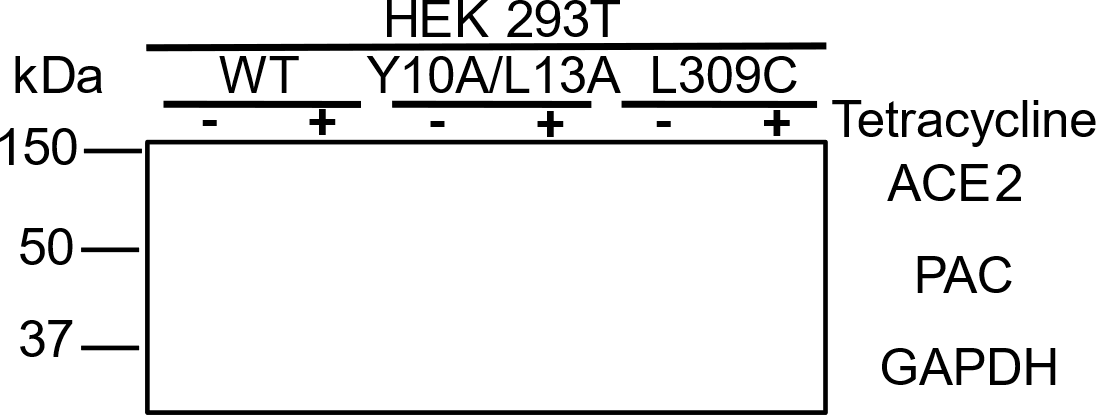

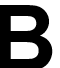

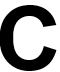

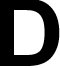

**Supplemental Figure 2. Validation of the PAC Y10A/L13A and L309C mutants.** A) Western blot of the PAC mutants. Representative image of three experiments. B) Representative immunofluorescence of the Y10A/L13A mutant showing reduced intracellular endosomal staining (labeled by EEA1) compared to WT. Scale bar: 5 μm. C) Representative I/V curve and D) current densities of acid-induced current of WT PAC and the channel “dead” mutant L309C. n = 10 cells from WT PAC inducible system. n = 13 cells from channel dead mutant. Bars represent mean ± SEM.

**Supplemental Figure 3. Validation of PAC expression and activity in Vero E6 cells.** A) Western blot of PAC protein. B) Representative I/V curve and C) current densities of acid-induced current in control and PAC-overexpressing Vero E6 cells. n = 3 cells. Bars represent mean ± SEM.

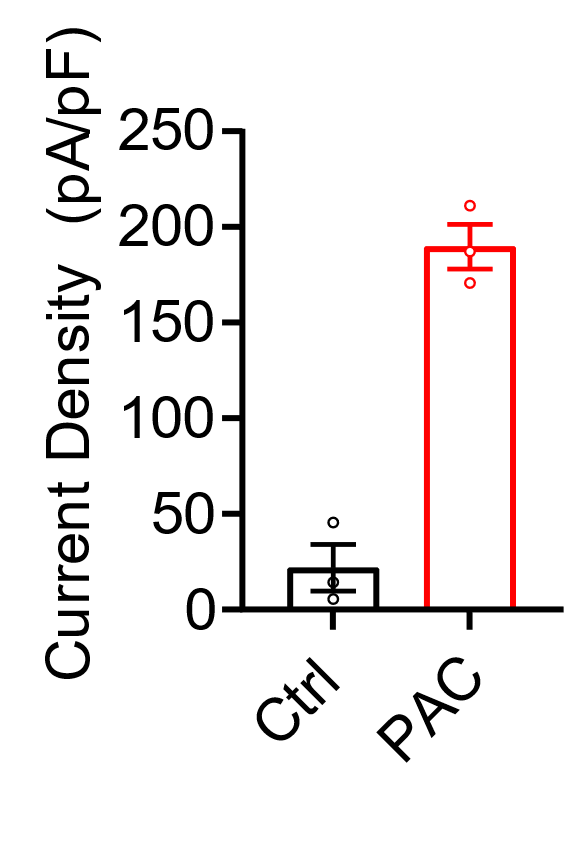

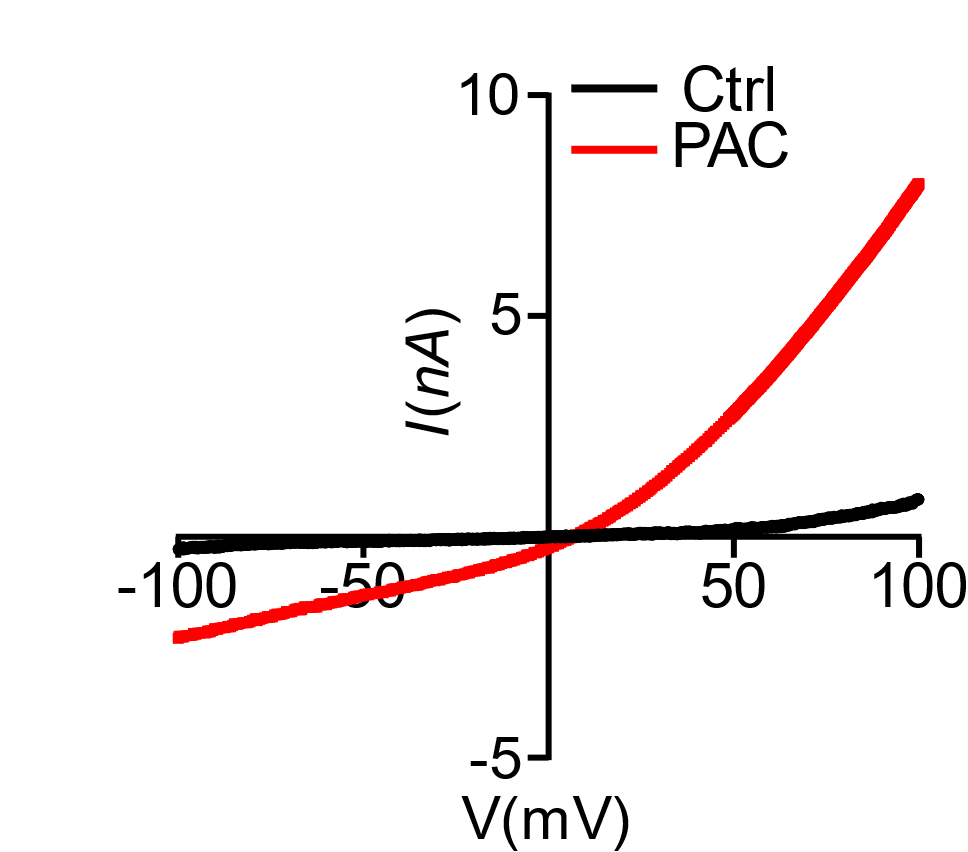

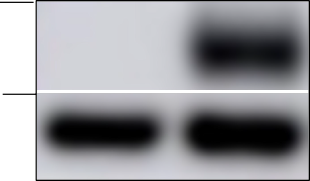

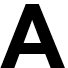

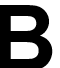

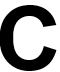

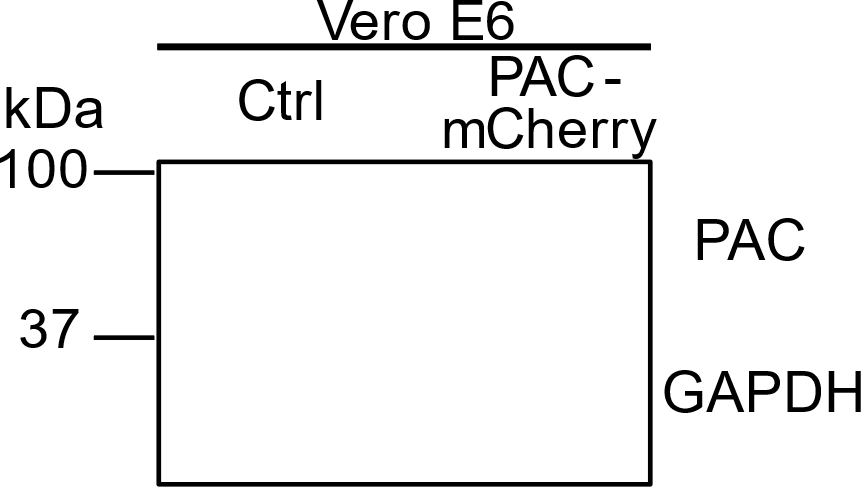

**Supplementary Table 1. List of reagents and plasmids used in this study.**

| Reagent or Resource | Source | Identifier | Reference |
| --- | --- | --- | --- |
| **Antibodies** | | |  |
| Mouse anti-PAC 4D12 | In house | N/A |  |
| Rat anti-hACE2 | Biolegend | Cat #375801; Clone A20069I |  |
| Goat anti-hACE2 | R&D systems | Cat #AF933 |  |
| Rabbit anti-TMPRSS2 | Abcam | ab109131; Clone EPR3862 |  |
| Mouse anti-GAPDH | Cell Signlaing Technology | Cat #97166 |  |
| Amersham ECL HRP-conjugated secondary antibody | Cytiva | Rabbit IgG, Rat IgG and Mouse IgG |  |
| Rabbit anti-EEA1 | Cell Signaling Technology | Cat #3288 |  |
| **Chemicals, peptides, and recombinant proteins** | | |  |
| Transferrin from human serum,Alexa Fluor633 conjugate | Thermo Fisher Scientific | Cat #T23362 |  |
| Transferrin from human serum, fluorescein conjugate | Thermo Fisher Scientific | Cat #T2871 |  |
| RIPA buffer | Sigma | Cat #R0278 |  |
| Intracellular pH Calibration Buffer Kit | Life Technologies | Cat #1787810 |  |
| Bright-Glo^TM^ Luciferase Assay System | Promega | Cat #E2620 |  |
| Tetracycline hydrochloride | Millipore Sigma | Cat #T4062 |  |
| Aloxistatin (E64-d) | Selleckchem | Cat #S7393 |  |
| **Experimental Cell Lines: Cell Models** | | |  |
| Human: Human embryonic kidney (HEK 293T) cells | ATCC | Cat #CRL-3216 |  |
| Human: Flip-InT-Rex 293 cells | Thermo Fisher Scientific | Cat #R78007 |  |
| Vero E6 | ATCC | CRL-1586 | A gift from Susan Weiss |
| **Plasmids** | | |  |
| pMD2.G(VSV-G) | Addgene | Plasmid #12259 | pMD2.G was a gift from Didier Trono |
| pMDLg/pRRE | Addgene | Plasmid #12251 | pMDLg/pRRE was a gift from Didier Trono |
| pRSV-Rev | Addgene | Plasmid #12253 | pRSV-Rev was a gift from Didier Trono |
| pHIV-Luc-ZsGreen | Addgene | Plasmid #39196 | pHIV-Luc-ZsGreen was a gift from Bryan Welm |
| HDM_SARS2_Spike_del21_D614G | Addgene | Plasmid #158762 | HDM_SARS2_Spike_del21_D614G was a gift from Jesse Bloom |
| pcDNA3.3_CoV1_D28 | Addgene | Plasmid #170447 | pcDNA3.3_CoV1_D28 was a gift from David Nemazee |
| pLENTI_hACE2_PURO | Addgene | Plasmid #155295 | pLENTI_hACE2_PURO was a gift from Raffaele De Francesco |
| pLEX307-TMPRSS2-blast | Addgene | Plasmid #158458 | pLEX307-TMPRSS2-blast was a gift from Alejandro Chavez & Sho Iketani |
| pcDNA3.3-SARS2-B.1.617.2 (Delta) | Addgene | Plasmid #172320 | pcDNA3.3-SARS2-B.1.617.2 (Delta) was a gift from David Nemazee |
| pcDNA3.3_SARS2_BQ.1.1 | Addgene | Plasmid #194493 | pcDNA3.3_SARS2_BQ.1.1 was d gift from David Nemazee |
| pcDNA3.3_SARS2_XBB | Addgene | Plasmid #194494 | pcDNA3.3_SARS2_XBB was a gift from David Nemazee |
| pcDNA3.3_SARS2_XBB.1.16 | Addgene | Plasmid #201189 | pcDNA3.3_SARS2_XBB.1.16 was a gift from David Nemazee |
| plenti_EF1aPACmCherry | Inhouse |  |  |
